# *ETHYLENE RESPONSE FACTOR 115* integrates jasmonate and cytokinin signaling machineries to repress adventitious rooting in *Arabidopsis*

**DOI:** 10.1101/2019.12.27.886796

**Authors:** Abdellah Lakehal, Asma Dob, Zahra Rahneshan, Ondřej Novák, Sacha Escamez, Sanaria Alallaq, Miroslav Strnad, Hannele Tuominen, Catherine Bellini

## Abstract

Jasmonate (JA), an oxylipin-derived phytohormone, plays crucial roles not only in plant immunity and defense against herbivorous insects but also in plant growth and developmental processes, including regeneration and organogenesis. However, the mechanistic basis of its mode of action and precise role in integrating other signaling cues are poorly understood. Here we provide genetic evidences that JA signaling acts in both NINJA-dependent and -independent modulation of the transcriptional activity of MYC transcription factors involved in the inhibition of adventitious root initiation (ARI). Our data show that NINJA-dependent JA signaling in pericycle cells blocks early events of ARI. Moreover, transcriptomic comparison of *ninja-1myc2-322B* double mutant (which produce extremely few ARs) and wild type seedlings identified a novel molecular network governed by the APETALA2/ETHYLENE RESPONSE FACTOR 115 (ERF115) transcription factor. We demonstrate that JA-induced *ERF115* activates the cytokinin signaling machinery and thereby represses ARI. Altogether, our results reveal a molecular network involving cooperative crosstalk between JA and CK machineries that inhibits ARI.

## INTRODUCTION

Jasmonate (JA), a stress-induced phytohormone, plays crucial roles in plant immunity and defense against herbivorous insects (Wasternack and Hause, 2013). It also participates in control of diverse developmental processes, including tissue regeneration and rhizotaxis (Wasternack and Hause, 2013; Lakehal et al., 2020). The isomer (+)-7-iso-JA-Ile (JA-Ile), the bioactive form of JA (Fonseca et al., 2009), is perceived by the F-box protein CORONATINE INSENSITIVE1 (COI1), which is an integral component of the Skp-Cullin-F-box (SCF) complex (Xie et al., 1998). The COI1 receptor fine-tunes the function of the JA transcriptional machinery in a simple manner. Briefly, in the resting state, marked by low JA-Ile contents, the transcriptional activity of a number of transcription factors, including the basic-Helix-loop-Helix MYC, is repressed by JASMONATE ZIM DOMAIN (JAZ) repressors through either physical interaction or recruitment of the general co-repressor TOPLESS (TPL) or TPL-related proteins (TPRs) (Chini et al., 2007; Thines et al., 2007; Yan et al., 2007). The adaptor NOVEL INTERACTOR OF JAZ (NINJA) mediates interaction of JAZs with TPL or TRPs (Pauwels et al., 2010). During activation, marked by accumulation of JA-Ile, JAZs form co-receptor complexes with COI1. This interaction is facilitated by JA-Ile, which acts as a molecular glue (Sheard et al., 2010). Formation of the co-receptor complexes triggers ubiquitylation and proteasome-dependent degradation of the targeted JAZs, thereby releasing the transcription factors to transcriptionally induce or repress their downstream target genes. Biochemical studies suggest that JAZ-dependent repression machinery can inhibit the transcriptional activity of different MYCs in different ways, depending on the JAZ protein involved (Chini et al., 2016). However, the biological roles of this multilayered regulation are unclear, largely because multiple *jaz* mutations may cause phenotypic deviations, but not single loss-of-function mutations (Campos et al., 2016; Guo et al., 2018).

JA signaling counteracts or cooperates with a number of hormonal and signaling cascades in the control of plant growth and development (Wasternack and Hause, 2013). We have previously shown that the COI1-dependent MYC2-mediated JA signaling inhibited the intact hypocotyl-derived ARI downstream of the auxin signaling machinery (Gutierrez et al., 2012) (Figure 1). Accordingly, in contrast to the *MYC2*-overexpressing line *35S:MYC2*, the loss-of-function mutant *myc2* produces more ARs than wild type plants, indicating that MYC2 plays an important role in inhibition of ARI downstream of auxin (Gutierrez et al., 2012). Recently, we also showed that the TIR1- and AFB2-dependent auxin signaling pathways promote ARI by negatively controlling JA content (Lakehal et al., 2019a). However, despite evidence of its central role in modulating ARI, the basis (genetic and mechanistic) and downstream targets of the MYC2-mediated JA signaling involved in this process remained unclear.

**Figure 1:**
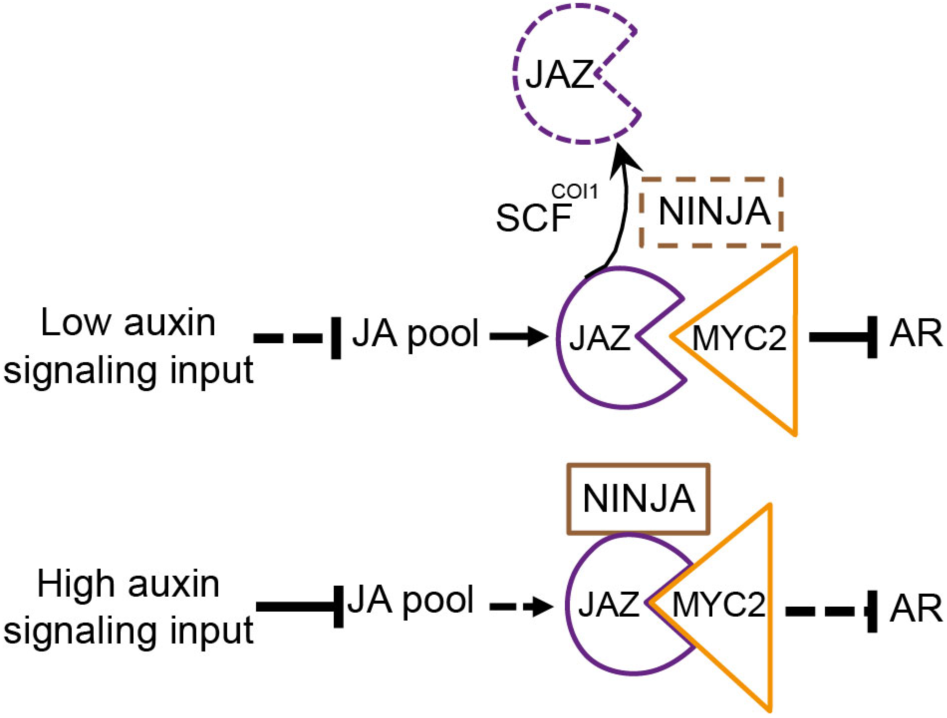
A genetic model for the action of JA signaling components during ARI. With a low auxin signaling input, the JA pool increases in the hypocotyl. This triggers degradation of the targeted JAZs, thereby releasing transcriptional activity of the MYC2, 3, 4 and inhibiting ARI. With a high auxin signaling input, the JA pool decreases in the hypocotyl, thereby repressing the MYC-mediated JA signaling machinery and increasing ARI (Gutierrez et al., 2012; Lakehal et al., 2019a).

Recently, Zhou and collaborators (Zhou et al., 2019) showed that two members of subgroup X of the *APETALA2/ETHYLENE RESPONSE FACTOR (ERF)* family (*ERF109 and ERF115*) promote root stem cell niche replenishment and tissue regeneration after excision, and their expression is directly controlled by MYC2-mediated JA signaling. The *ERF115* transcription factor and its two closest homologs, *ERF114 (*also known as *ERF BUD ENHANCER* (*EBE*) and *ERF113* (also known as *RELATED to AP2.6L*, *RAP2.6L)* have been shown to control various regenerative processes, such as callus formation, tissue repair, root stem cell niche maintenance and root growth (Che et al., 2006; Nakano et al., 2006; Asahina et al., 2011; Mehrnia et al., 2013; Heyman et al., 2016; Ikeuchi et al., 2018; Kong et al., 2018; Yang et al., 2018). The three genes are rapidly induced by mechanical wounding (Ikeuchi et al., 2017), suggesting that they play an important role in connecting the stress-induced JA signaling machinery with other signaling cascades in provision of correct cell-fate and/or developmental inputs for organogenesis processes. However, how these genes coordinate and integrate the stress-induced hormonal pathways to ensure these multifunctionalities is still largely unclear. Here we provide evidence that the JA signaling machinery inhibits ARI in both NINJA-dependent and - independent manners, and the JA-induced *ERF115* transcription factor inhibits this process in a CK-dependent manner, suggesting that CKs act downstream of JA in ARI inhibition.

## RESULTS

### NINJA-dependent and -independent JA signaling repress ARI

To better understand the role of JA signaling during intact hypocotyl-derived ARI (Figure 1), we first analyzed the AR phenotype of multiple *jaz* mutants, under previously described conditions (Sorin et al., 2005; Gutierrez et al., 2009; Gutierrez et al., 2012). The quadruple loss-of-function mutant *jaz7jaz8jaz10jaz13* (Thireault et al., 2015) had the same phenotype as the wild type, whereas the quintuple mutant *jazQ (jaz1jaz3jaz4jaz9jaz10)* (Campos et al., 2016) produced slightly fewer ARs than wild type plants (Figure 2A). These data confirm the high functional redundancy of the 13 *JAZ* genes in the Arabidopsis genome (Chini et al., 2007; Thines et al., 2007; Yan et al., 2007; Thireault et al., 2015; Chini et al., 2016), which complicates characterization of their specificity. Therefore, we analyzed the phenotype of the gain-of-function mutant *myc2-322B*, which harbors a point mutation in the transcriptional activation domain (TAD) that changes Glutamate 165 to Lysine. This prevents MYC2’s interaction with most JAZ repressor proteins, resulting in almost constitutive MYC2 signaling (Gasperini et al., 2015). We found that *myc2-322B* produced slightly fewer AR than wild type plants (Figure 2B), in accordance with our previous findings that the loss-of-function mutant *myc2* and overexpressing line *35S:MYC2* respectively produced more and less ARs than wild type counterparts (Gutierrez et al., 2012). We also analyzed the AR phenotype associated with two loss-of-function *ninja (ninja-1* and *ninja-2)* alleles (Acosta et al., 2013), because the NINJA adaptor is a central hub in the transcriptional repression machinery that inactivates MYC transcription factors (Pauwels et al., 2010) (Figure 1). *Ninja-1* and *ninja-2* mutants produced significantly fewer ARs than wild type plants (Figure 2B), but their phenotypic deviation is weak, presumably due to presence of a NINJA-independent pathway that continues to repress MYCs and thus allows ARI. Because MYC2 acts additively with MYC3 and MYC4 in the inhibition of ARI (Gutierrez et al., 2012), we hypothesized that removing NINJA in a *myc2-322B* background might abolish the remaining NINJA-dependent repression and hence release activity of the three MYCs. De-repression of these transcription factors would then result in constitutively enhanced MYC-mediated JA signaling and block the ARI process. To test this hypothesis, we analyzed the AR phenotype of two independent double mutants: *ninja-1myc2-322B* and *ninja-2myc2-322B* (Gasperini et al., 2015). We found that ARI was almost completely inhibited in both double mutants, confirming the inhibitory effect of JA (Figure 2B-E). As expected, the double mutants had shorter primary roots (PRs) than wild type plants, due to the inhibitory effect of JA signaling on PR growth (Staswick et al., 1992) and fewer lateral roots (LRs; Supplemental Figure 1A,B), but the LR density was not affected (Figure 2C). To get further genetic evidence, we also analyzed the AR phenotype of the gain-of-function mutant *atr2D,* which harbors a point mutation in the JAZ interaction domain (JID) of the MYC3 protein (Smolen et al., 2002) that prevents its interaction with a subset of JAZ repressors (Zhang et al., 2015). Notably, there was no significant difference in AR numbers of *atr2D* mutants and wild type plants, but the *ninja-1atr2D* double mutant produced far fewer ARs (Figure 2B), confirming the *atr2D* mutation’s additive effect and the role of MYC3 in the control of AR formation. Collectively, these results genetically confirm the importance of the NINJA-dependent and -independent pathways in the control of AR initiation.

**Figure 2:**
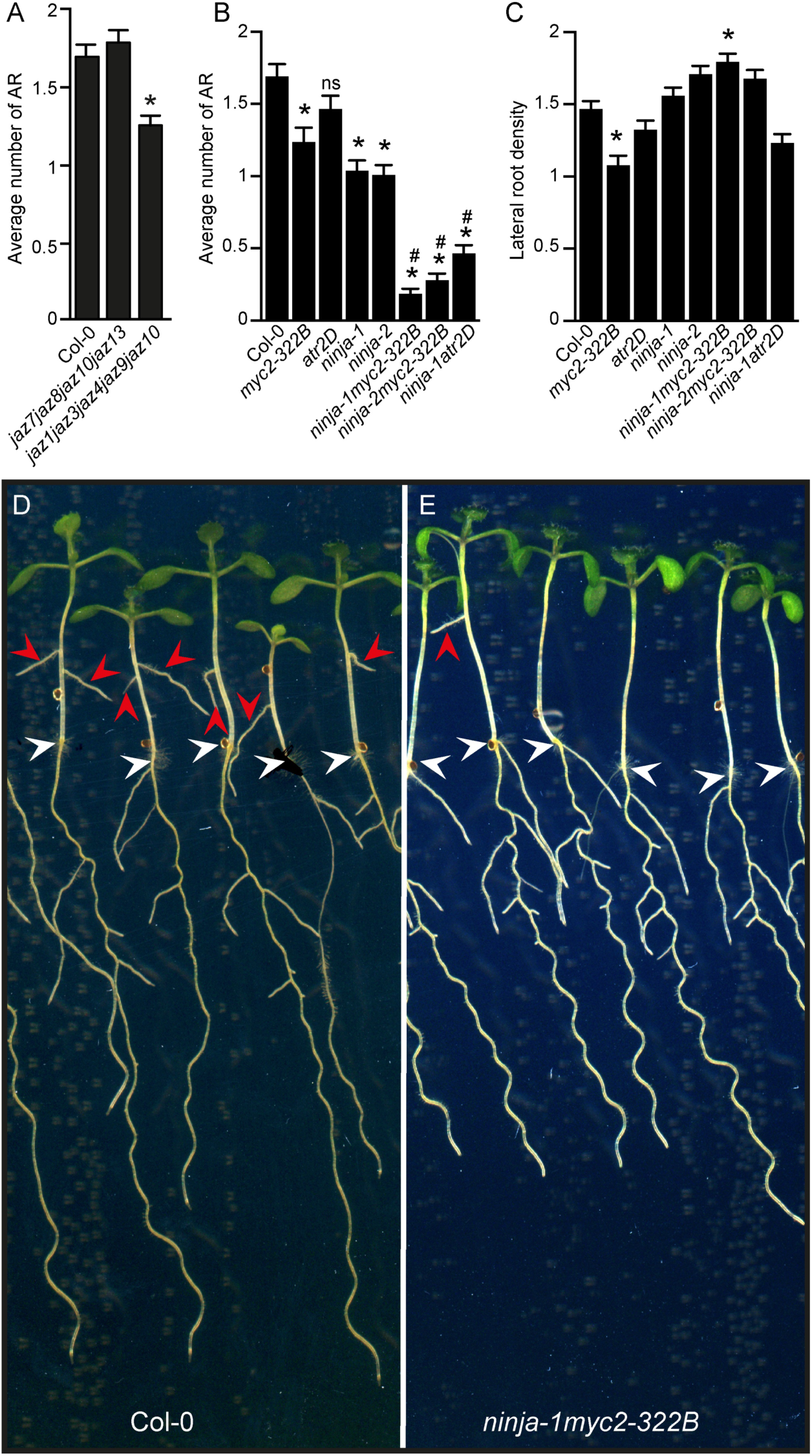
JA signaling inhibits ARI in NINJA-dependent and -independent manners. (A) Average number of ARs observed in indicated multiple *jaz* mutants and wild-type (Col-0) seedlings. Data are pooled and averaged numbers observed in three biological replicates of at least 40 seedlings. One-way ANOVA combined with Tukey’s multiple comparison post-tests showed that the *jaz1jaz3jaz4jaz9jaz10* produced significantly more ARs than wild-type plants. Error bars indicate ± SEM (n ≥ 40; P < 0.05). (B) Average number of ARs produced by JA signaling mutants. A non-parametric Kruskal-Wallis test followed by Dunn’s multiple comparison test indicated that mutations in the *MYC2* or *NINJA* genes result in significant differences in AR number, relative to wild-type numbers. Error bars indicate ± SEM (n ≥ 40; P < 0.02). Values marked with asterisks significantly differ from corresponding wild-type values and those marked with hash signs significantly differ from values obtained for the single *ninja-1 or ninja-2* mutants. (C) Lateral root density of JA signaling mutants and wild-type seedlings grown in AR phenotyping conditions. One-way ANOVA combined with Tukey’s multiple comparison post-test indicated that the *myc2-322B* and *ninja-1myc2-322B* mutants had slightly lower and slightly higher than wild-type LR densities, respectively. Error bars indicate ± SEM (n ≥ 40; P < 0.05). (D) to (E) Representative photos of (D) wild-type and (E) *ninja-1myc2-322B* double mutant seedlings. Scale bars represent 6 mm. Arrowheads indicate hypocotyl-root junctions (white) or ARs (red).

### *NINJA* and *MYC2* are expressed in the etiolated hypocotyl

To examine spatiotemporal expression patterns of the *NINJA* and *MYC2* genes during early ARI events, we used seedlings harboring *pNINJA:GUS* or *pMYC2:GUS* transcriptional fusions (Gasperini et al., 2015). The seedlings were grown in ARI-inducing conditions in the dark and sampled for *pNINJA:GUS* or *pMYC2:GUS* expression analysis at T0, just before some of the etiolated seedlings were exposed to light. Further samples were collected at T9L and T24L (after 9 and 24 h growth in long-day conditions, respectively), while controls were sampled at T9D and T24D (after a further 9 and 24 h growth in the dark, respectively). The two promoters were shown to be constitutively active in all the organs at all time points, although *MYC2* promoter activity declined in the cotyledons over time (Figure 3A to E). These data indicate that *NINJA* and *MYC2* genes have overlapping expression domains in the hypocotyl.

**Figure 3:**
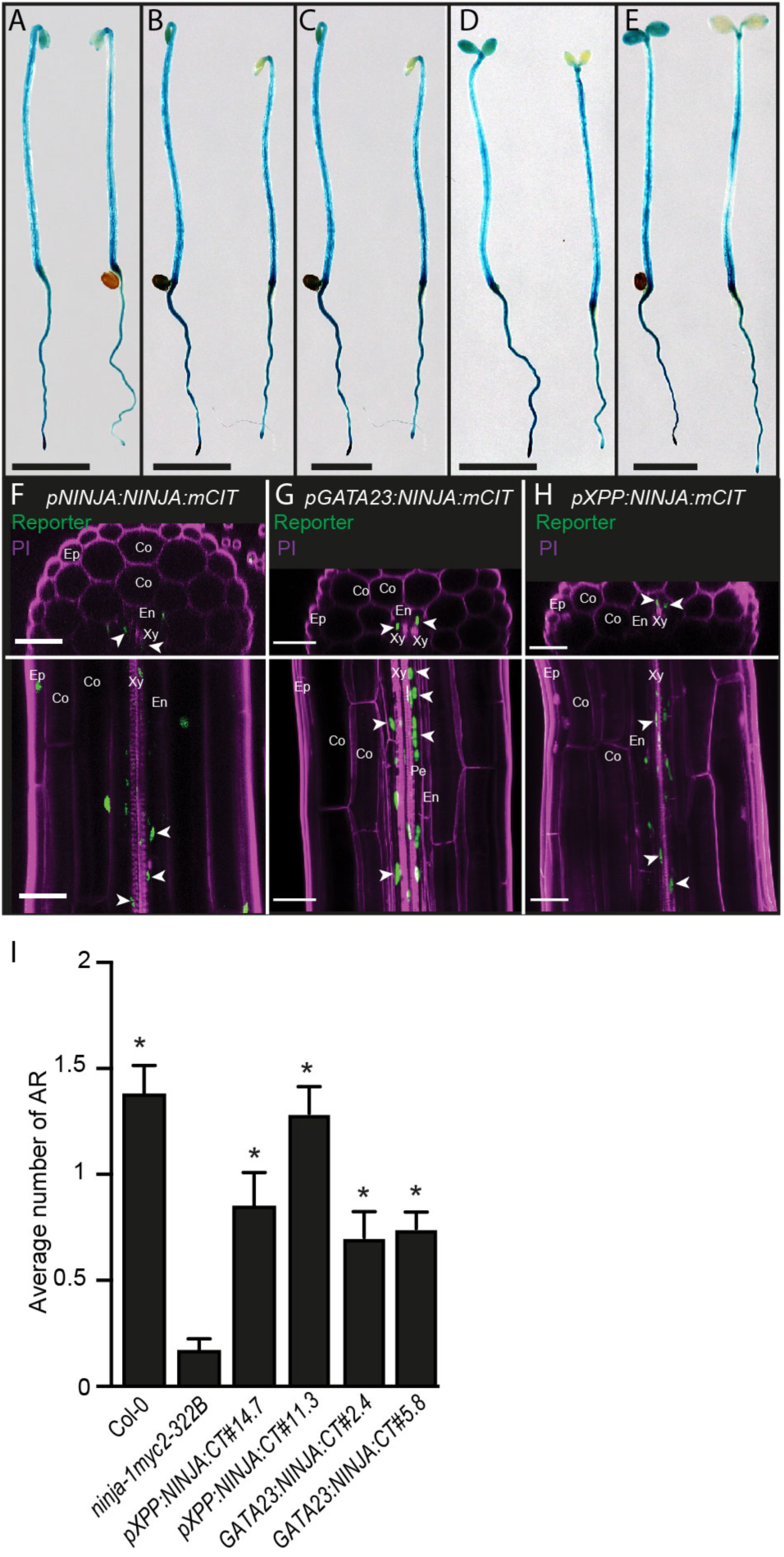
NINJA-dependent JA signaling inhibit ARI in pericycle cells. (A) to (E) Spatiotemporal activity patterns of the *NINJA* and *MYC2* promoters, left and right, respectively in each panel. Seedlings expressing the *pNINJA:GUSplus* or *pMYC2:GUSplus* constructs were grown in the dark until their hypocotyls were 6-7 mm long (T0) (A) then either kept in the dark for 9 h (T9D) (B) and 24 h (T24D) (C) or transferred to the light for 9 h (T9L) (D) or 24 h (T24L) (E). Scale bars represent 6 mm. (F) to (H) Representative images of etiolated hypocotyls expressing *pNINJA:NINJA:mCITRINE* (F), *pGATA23:NINJA:mCITRINE* (G), and *pXPP:NINJA:mCITRINE* (H) of seedlings grown in darkness until their hypocotyls were 6-7 mm long. The cell walls were counterstained magenta with propidium iodide (PI). Orthogonal views from epidermis to vasculature are shown in the top panels. Z-projections of the hypocotyl volume around the vasculature are shown in the bottom panels. The following cell types can be distinguished: Epidermis (Ep), Cortex (Co), Endodermis (En), Pericycle (Pe) and Xylem (Xy). In orthogonal views, the two protoxylem elements allow deduction of the direction of the xylem axis and thus the position of the xylem-pole pericycle. Arrowheads indicate signals in xylem-pole pericycle cells in green. (I) Average numbers of ARs produced by the *ninja-1myc2-322B* double mutant, two independent transgenic lines expressing *pXPP:NINJA:mCITRINE/ninja-1myc2-322B* or *pGATA23:NINJA:mCITRINE/ninja-1myc2-322B* and wild-type (Col-0) seedlings. A non-parametric Kruskal-Wallis test followed by Dunn’s multiple comparison post-test indicated that *pXPP:NINJA:mCITRINE/ninja-1myc2-322B* (#14.7 and #11.3) and *pGATA23:NINJA:mCITRINE/ninja-1myc2-322B* (#2.4 and #5.8) produced significantly more ARs than the *ninja1myc2-322B* double mutant. Error bars indicate ± SEM (n ≥ 30; P < 0.006).

### Expressing NINJA in xylem-pole pericycle cells is sufficient to counter JA’s negative effect during ARI

We confirmed that the NINJA protein was broadly expressed in the hypocotyl, including the xylem-pole pericycle (xpp) cells (Figure 3F) where ARs are initiated (Sorin et al., 2005; Sukumar et al., 2013). We then assessed whether re-activating the NINJA-dependent JA repression machinery in those cells would be sufficient to restore ARI in the *ninja1-myc2-322B* double mutant. For this, we produced translational fusions of NINJA with the mCITRINE reporter driven by two xpp cell-specific promoters, *GATA23* (De Rybel et al., 2010) and *XPP* (Andersen et al., 2018). The p*GATA23:NINJA:mCITRINE* or p*XPP:NINJA:mCITRINE* constructs were introduced into the *ninja-1myc2-322B* double mutant, and we confirmed that the NINJA:mCITRINE protein was specifically present in the hypocotyl xpp cells (Figure 3G and 3H). We analyzed the AR phenotype of two independent lines carrying each construct and showed that in both cases the effect of the *ninja-1* mutation was complemented (Figure 3I). These results suggest that expressing NINJA in xpp cells is sufficient to de-repress ARI, and that NINJA-dependent JA signaling acts in early stages of ARI.

### Transcriptomic insights into JA’s role in ARI

To get mechanistic insights into how JA signaling reprograms the transcriptional machinery during ARI, we compared transcriptomes of *ninja-1myc2-322B* double mutant and wild-type hypocotyls at three time points: T0, T9 and T24 (Figure 4A). In T0 samples we detected 530 differentially expressed genes (DEGs), of which 462 were upregulated and 68 downregulated in the *ninja-1myc2-322B* double mutant. We detected 671 DEGs at T9, 453 upregulated and 218 downregulated, and 579 at T24, 388 upregulated and 191 downregulated (Figure 4B, Supplemental Figure 2 and Supplemental Table 1).

**Figure 4:**
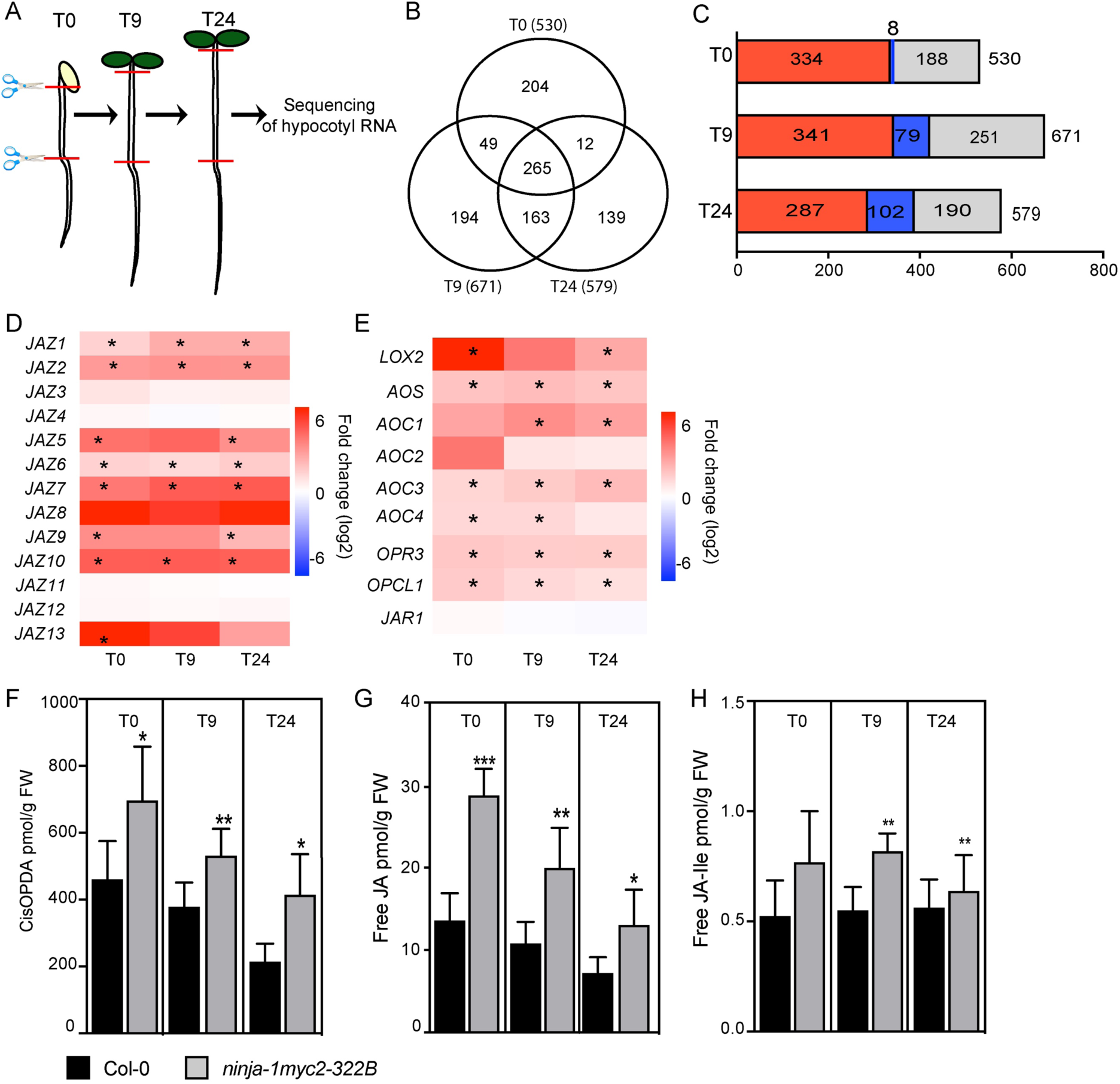
RNA-Seq revealed several DEGs between the *ninja-1myc2-322B* double mutant and wild-type seedlings. (A) Schematic representation of the RNA-Seq experiment. Total RNA was extracted from hypocotyls of *ninja-1myc2-322B* double mutant and wild-type seedlings grown in the dark until their hypocotyls were 6-7 mm long (T0), and after their transfer to the light for 9 h (T9) or 24 h (T24). (B) Venn diagram summarizing the DEGs between *ninja-1myc2-322B* double mutant and wild-type seedlings. (C) Enrichment of G-box (CACGTG, CACATG) or G-box-like (AACGTG, CATGTG CACGCG or CACGAG) motifs in the DEGs. Colors indicate upregulated genes (red) or downregulated genes (blue) containing at least one of the motifs. The gray color indicates the remaining DEGs, containing none of the mentioned motifs. (D) Heatmap of expression of the 13 *JAZ* genes. The map is based on fold-differences (log_2_) in transcript abundance (based on RNA-Seq data) in *ninja-1myc2-322B* double mutant samples relative to the abundance in wild-type samples. Colors indicate upregulated genes (red) and downregulated genes (blue) in *ninja-1myc2-322B* double mutant relative to expression levels in wild-type seedlings. Values marked with asterisks are statistically significant. (E) Heatmap of expression selected JA biosynthesis genes. The map is based on fold-differences (log_2_) in transcript abundance (based on RNA-Seq data) in *ninja-1myc2-322B* double mutant samples relative to the abundance in wild-type samples. Colors indicate upregulated genes (red) and downregulated genes (blue) in *ninja-1myc2-322B* relative to wild-type expression levels. Values marked with asterisks are statistically significant. (F) to (H) Endogenous jasmonate contents. (F) *cis*-OPDA, (G) free JA and (H) JA-Ile contents of hypocotyls of *ninja-1myc2-322B* and wild-type seedlings grown in the dark until their hypocotyls were 6 mm long (T0) and after their transfer to the light for 9 h (T9) and 24 h (T24). Asterisks indicate statistically significant differences between the mutant lines and wild-type plants according to ANOVA analysis (*, **, and *** indicate p values of 0.05 > p > 0.01, 0.01 > p > 0.001, and p < 0.001, respectively). Error bars indicate ± SD of six biological replicates.

### The *ninja-1myc2-322B* double mutant has a constitutive JA response signature

MYC transcription factors recognize and bind to hexameric *cis*-regulatory G-box motifs (CACGTG or CACATG), and MYC2 binds to G box-like motifs (AACGTG, CATGTG, CACGAG, CACATG, CACGCG) with differing affinities (Godoy et al., 2011). To get an overview of possible direct targets of MYCs among the DEGs, we searched for these motifs in the 1 kb regions upstream of their ATG translation start codons. We found that DEGs’ promoters are highly enriched with MYC binding sites, suggesting that they include potential direct targets of MYC. At T0, T9 and T24, 64% of 520 DEGs (342: 334 upregulated and 8 downregulated), 62% of 671 DEGs (420: 341 upregulated and 79 downregulated), and 67% of 579 DEGs (389: 287 upregulated and 102 downregulated) respectively contained at least one of the six motifs (Figure 4C).

Most of the *JAZ* genes, which are early JA-responsive genes, were highly upregulated in the *ninja-1myc2-322B* double mutant at all sampling time points (Figure 4D), confirming the presence of enhanced, constitutive JA signaling. Accordingly, several genes involved in JA biosynthesis, such as *LIPOXYGENASE 2 (LOX2), ALLENE OXIDE SYNTHASE (AOS), ALLENE OXIDE CYCLASE1 (AOC1), AOC3, AOC4, OXOPHYTODIENOATE-REDUCTASE3 (OPR3) and OPC-8:0 COA LIGASE1 (OPCL1)* were upregulated in the double mutant *ninja-1myc2-322B* (Figure 4E). The biological relevance of this upregulation of gene expression was confirmed by findings that levels of the JA precursor *cis*-12-oxo-phytodienoic acid (*cis*-OPDA), JA and JA-Ile were higher in the double mutant than in wild-type controls at all time points, except that JA-Ile contents did not significantly differ at T0 (Figure 4F-H). These data highlight a feedforward loop that amplifies the response to JA signaling by enhancing JA biosynthesis.

### JA signaling controls expression of *ERF113*, *ERF114* and *ERF115* transcription factors

The candidate transcription factor potentiel targets of MYC2 we detected included three closely related members of subgroup X of the *ERF* family (*ERF113*, *ERF114* and *ERF115*) (Figure 5A,B). Analysis by qRT-PCR confirmed that these three genes were all upregulated in the hypocotyl of the *ninja-1myc2-322B* double mutant, except *ERF113* at T0 (Figure 5C). These genes have known involvement in a number of organogenesis and regeneration processes (Heyman et al., 2018). To address their role in ARI, we analyzed the AR phenotypes of available single loss of *ERF113* or *ERF115* function mutants (*rap2.6l-1* and *erf115,* respectively) and observed no significant difference in this respect between them and wild-type controls (Figure 6A). As no loss-of-function T-DNA line for *ERF114* was available, we used CRISPR-Cas9 technology to delete a ca. 40 bp genomic fragment in the first exon of the *ERF114* gene in the *rap2.6l-1* and the *erf115* backgrounds to obtain *rap2.6l-1erf114C* and *erf115erf114C* double mutants, respectively (Supplemental Figure 3). Other multiple mutants were obtained by genetic crosses. Only the triple mutant *rap2.6l-1erf114Cerf115* produced significantly more ARs than wild-type controls (Figure 6A), indicating that *ERF113, ERF114* and *ERF115* act redundantly in the control of ARI.

**Figure 5:**
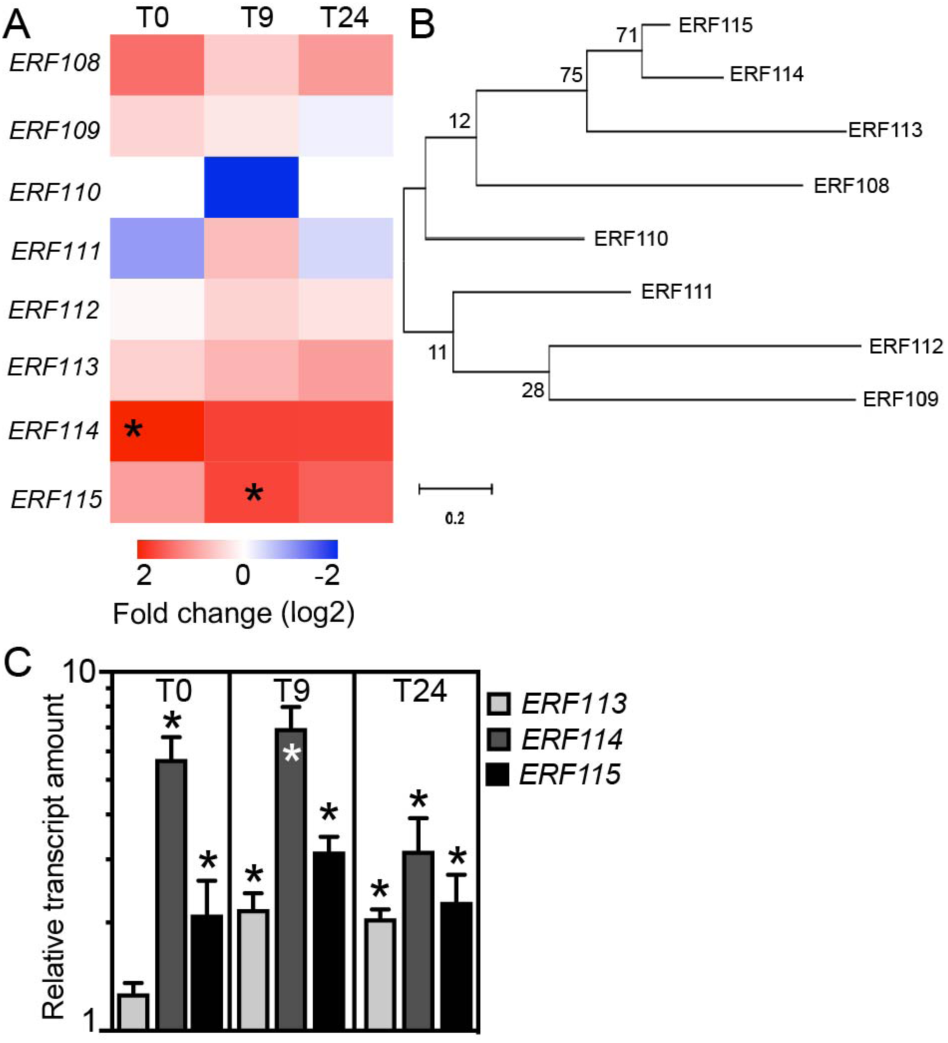
*ERF113*, *ERF114* and *ERF115* are induced by JA signaling. (A) Heatmap of expression of the subgroup X *ERF family* members. The map is based on fold-differences (log_2_) in transcript abundance (based on RNA-Seq data) in *ninja-1myc2-322B* double mutant samples relative to the abundance in wild-type samples. Colors indicate upregulated genes (red) or downregulated genes (blue) in *ninja-1myc2-322B* relative to wild type expression levels. Values marked with asterisks are statistically significant. (B) Phylogenetic tree of subgroup X of the AP2/ERF protein family derived from protein sequence alignment by the maximum likelihood method using MEGA X software (Kumar et al., 2018). (C) Validation by qRT-PCR of mutation-induced shifts in *ERF113*, *ERF114* and *ERF115* expression profiles in the *ninja-1myc2-322B* double mutant (abundance of transcripts, in log10 scale, at indicated time points relative to their abundance in wild-type seedlings, which was arbitrarily set to 1). Error bars indicate ±SE obtained from three independent technical replicates. Asterisks mark significance differences between the genotypes according to a *t*-test (P < 0.001, n = 3). The experiment was repeated twice with independent biological replicates and gave similar results.

**Figure 6:**
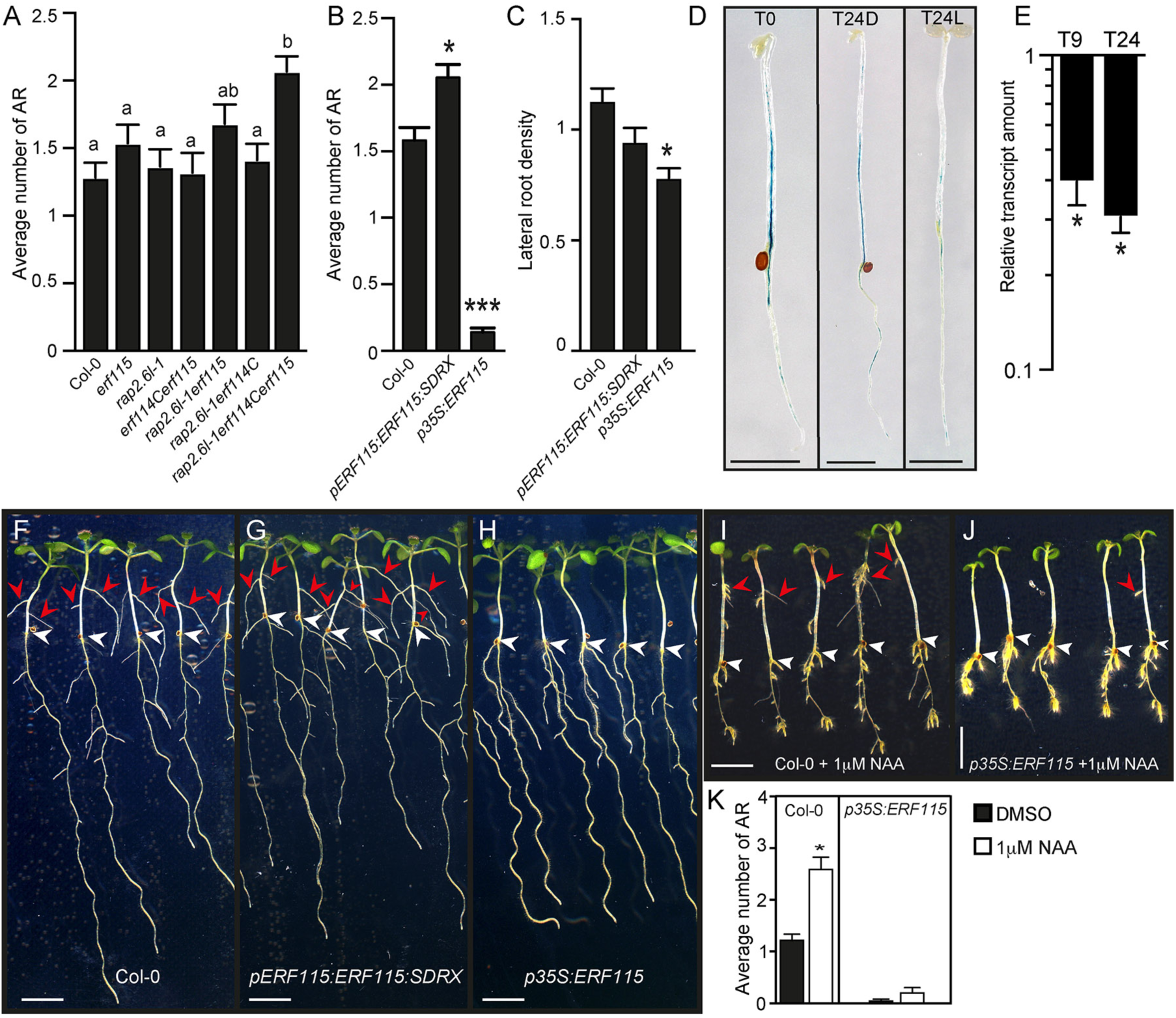
The *ERF115* gene is an inhibitor of ARI. (A) Average numbers of AR produced by *erf* mutants and wild-type seedlings. One-way ANOVA combined with Tukey’s multiple comparison post-test indicated that only the triple mutant *rap2-6lerf114Cerf115* significantly differed in this respect from wild-type (Col-0) plants. Error bars indicate ± SEM (n ≥ 40, P < 0.001). (B) Average numbers of ARs produced by *35S:ERF115* and *pERF115:ERF115:SRDX* relative to numbers produced by wild-type plants. Data from two independent biological replicates, each of at least 40 seedlings, were pooled and averaged. A non-parametric Kruskal-Wallis test followed by Dunn’s multiple comparison test indicated that numbers of ARs produced by the transgenic and wild-type plants significantly differed. Error bars indicate ± SEM (n ≥ 40, P < 0.02). (C) LR density of *35S:ERF115* and *pERF115:ERF115:SRDX* lines and wild type plants in AR phenotyping conditions. *35S:ERF115* mutants had significantly lower LR density than wild-type plants according to one-way ANOVA followed by Tukey’s multiple comparison test. (D) Spatiotemporal activity pattern of the *ERF115* promoter, as shown by seedlings expressing the *pERF115:GUS* construct grown in the dark until their hypocotyls were 6-7 mm long (T0), and 24 h (T24L) after either transfer to the light or further growth in the dark (T24D). Scale bars represent 6 mm. (E) Validation by qRT-PCR of *ERF115* expression patterns in wild-type plants. Presented gene expression values are relative (in log10 scale) to the expression at T0, for which the value was arbitrarily set to 1. Error bars indicate ±SE obtained from three independent technical replicates. A *t*-test indicated that values indicated by an Asterisks indicate values that significantly differ from the T0 values (P < 0.001, n = 3). The experiment was repeated twice with independent biological replicates and gave similar results. (F) to (H) Representative photos of (F) wild-type, (G) *35S:ERF115*, and (H) *pERF115:ERF115:SRDX* seedlings. (I) to (J) Representative photos of wild-type and *35S:ERF115* seedlings grown in the dark until their hypocotyls were 6-7 mm long, then transferred to fresh medium containing either mock solution or 1 µM naphthaleneacetic acid (NAA) for seven more days under long-day conditions to induce ARs. Arrowheads indicate hypocotyl-root junctions (white) or ARs (red). Scale bars represent 6 mm. (K) Average numbers of ARs produced by wild-type and *35S:ERF115* plants in response to NAA. Wild-type seedlings produced significantly more ARs after NAA treatment than after mock-treatment according to a Mann-Whitney test (n ≥ 40, *P* < 0.0001), but NAA treatment had no significant effect on AR production by *35S:ERF115* plants. Error bars indicate ± SEM.

### *ERF115* represses hypocotyl-derived AR initiation downstream of auxin

Previous findings that *ERF115*’s expression is directly controlled by MYC2 and it plays major roles in root regeneration and stem cell replenishment (Heyman et al., 2013; Heyman et al., 2016; Zhou et al., 2019) prompted us to address its function during ARI. First, to overcome potential functional redundancy with other members of the family, we analyzed the *pERF115:ERF115:SRDX* line, which expresses a dominant negative variant of ERF115 (because the *ERF115* coding sequence fused to the ethylene-responsive element binding factor-associated amphiphilic repression (EAR) domain is driven by the *ERF115* promoter to ensure repression in the native expression domain (Heyman et al., 2013). The *pERF115:ERF115:SRDX* line produced significantly more ARs than wild-type controls, but was very similar to the *rap2.6l-1erf114Cerf115* triple mutant (Figure 6A,B). Although we cannot exclude a potential contribution of other *ERF* genes, these findings suggest that *ERF113*, *ERF114* and *ERF115* are the main transcription factors involved in ARI. Interestingly, the overexpressing line *35S:ERF115* developed extremely few ARs (Figure 6B) but had only slightly lower LR density than wild-type plants (Figure 6C and Supplemental Figure 4C,D). Thus, it phenocopied the *ninja-1myc2-322B* double mutant and confirmed that *ERF115* is an ARI repressor. We also characterized *ERF115*’s expression pattern during early ARI events using lines harboring the transcriptional fusion *pERF115:GUS*(Heyman et al., 2013). At T0, GUS staining was mainly detected in vascular tissues of the hypocotyl, and to a lesser extent in the root (Figure 6D). Exposing the seedlings to light for 24 h dramatically decreased the GUS signal (Figure 6D), suggesting that the *ERF115* gene is expressed in vascular tissue and its expression is negatively regulated by light, which we confirmed by qRT-PCR (Figure 6E). As JA acts downstream of auxin signaling in ARI inhibition (Gutierrez et al., 2012; Lakehal et al., 2019a), we hypothesized that the *35S:ERF115* line could be insensitive to exogenously applied auxin. To test this hypothesis, we treated *35S:ERF115*-expressing and wild-type pre-etiolated seedlings with the synthetic auxin naphthaleneacetic acid (NAA), and found that 1 μM NAA significantly enhanced AR development in the wild-type seedlings, but did not affect the *35S:ERF115*-expressing seedlings (Figure 6I-K). These data suggest that auxin cannot bypass the inhibitory effect of *ERF115* during ARI. Notably, the PR and LRs of the *35S:ERF115*-expressing seedlings were as sensitive as the wild-type roots to NAA (Figure 6I,J). These data suggest that *ERF115* specifically activates and/or cooperates with other negative regulator(s) of ARI downstream of auxin signaling.

### *ERF115*-mediated ARI repression requires cytokinins (CKs)

CKs, in balance with auxin, are known to promote shoot and callus formation but inhibit root growth and AR formation (Lakehal and Bellini, 2018; Ikeuchi et al., 2019), raising the possibility that modulation of the CK machinery by *ERF115* is involved in this multifunctionality. We confirmed the negative role of CKs in control of ARI as exogenously applied 6-benzyladenine (6-BA) inhibited the process in a dose-dependent manner (Figure 7A). We then analyzed the CK-deficient triple loss*-*of*-*function mutant *ipt3ipt5ipt7* that lacks three important ATP/ADP ISOPENTENYLTRANSFERASES catalyzing a rate-limiting step in *de novo* CK biosynthesis (Miyawaki et al., 2006), and a line overexpressing *CYOKININ OXIDASE1* (*35S:CKX1)*, which is also deficient in CKs due to their enhanced degradation(Werner et al., 2003). Notably, both the triple loss*-*of*-*function mutant *ipt3ipt5ipt7* and the *35S:CKX1*-expressing line produced significantly more ARs than wild-type controls (Figure 7B,C). Similarly, the *arr1-3arr11-2* double mutant and *arr1-3arr11-2arr12-1* triple mutant, which lack the key type-B transcription factors ARR1, ARR11 and ARR12 involved in CK signaling, produced significantly more ARs than wild-type plants (Figure 7D). These data genetically confirmed that CKs are repressors of ARI.

To test the hypothesis that *ERF115* inhibits ARI through CKs, we quantified relative amounts of transcripts of two CK-responsive genes, *ARR5* and *ARR7*, in etiolated hypocotyls of the overexpressing line *35S:ERF115* and wild-type controls at T0 and T24. Interestingly, at T0 *ARR7* was upregulated, and at T24 both *ARR5* and *ARR7* were upregulated in the *35S:ERF115* line (Figure 7E). These findings suggest that CK responses are enhanced in hypocotyls of *35S:ERF115* plants, and to explore possible causes we quantified active CK bases at T0, T9 and T24. At T0, isopentyladenine (iP), *trans*-Zeatin (*t*Z) and *cis*-Zeatin (*c*Z) contents of *35S:ERF115* and wild-type plants did not significantly differ (Figure 7F). However, at T9, *35S:ERF115* plants had significantly higher iP, *t*Z and *c*Z contents, and at T24 significantly higher iP and *c*Z contents than wild-type controls (Figure 7F). These data suggest that the *ERF115* inhibits ARI by modulating the CK pool. To test this hypothesis, we generated a *35S:ERF115ipt3ipt5ipt7* multiple mutant and a line overexpressing both *35S:ERF115* and *35S:CKX1* to deplete the CK pool in a *35S:ERF115* background, and confirmed that this was sufficient to restore ARI to wild-type levels in the *35S:ERF115* line (Figure 7G). These data indicate that *ERF115* inhibition of ARI is mediated by CKs. Interestingly, our transcriptomic data showed that several *LONELY GUY* (*LOG*) genes, which control a rate-limiting step in CK biosynthesis (Kuroha et al., 2009), were slightly upregulated, while several *CKX* genes were slightly downregulated, in the *ninja-1myc2-322B* double mutant (Supplemental Figure 5A,B). Altogether, our results strongly suggest that JA inhibits ARI by modulating CK homeostasis through the action of *ERF115*

**Figure 7:**
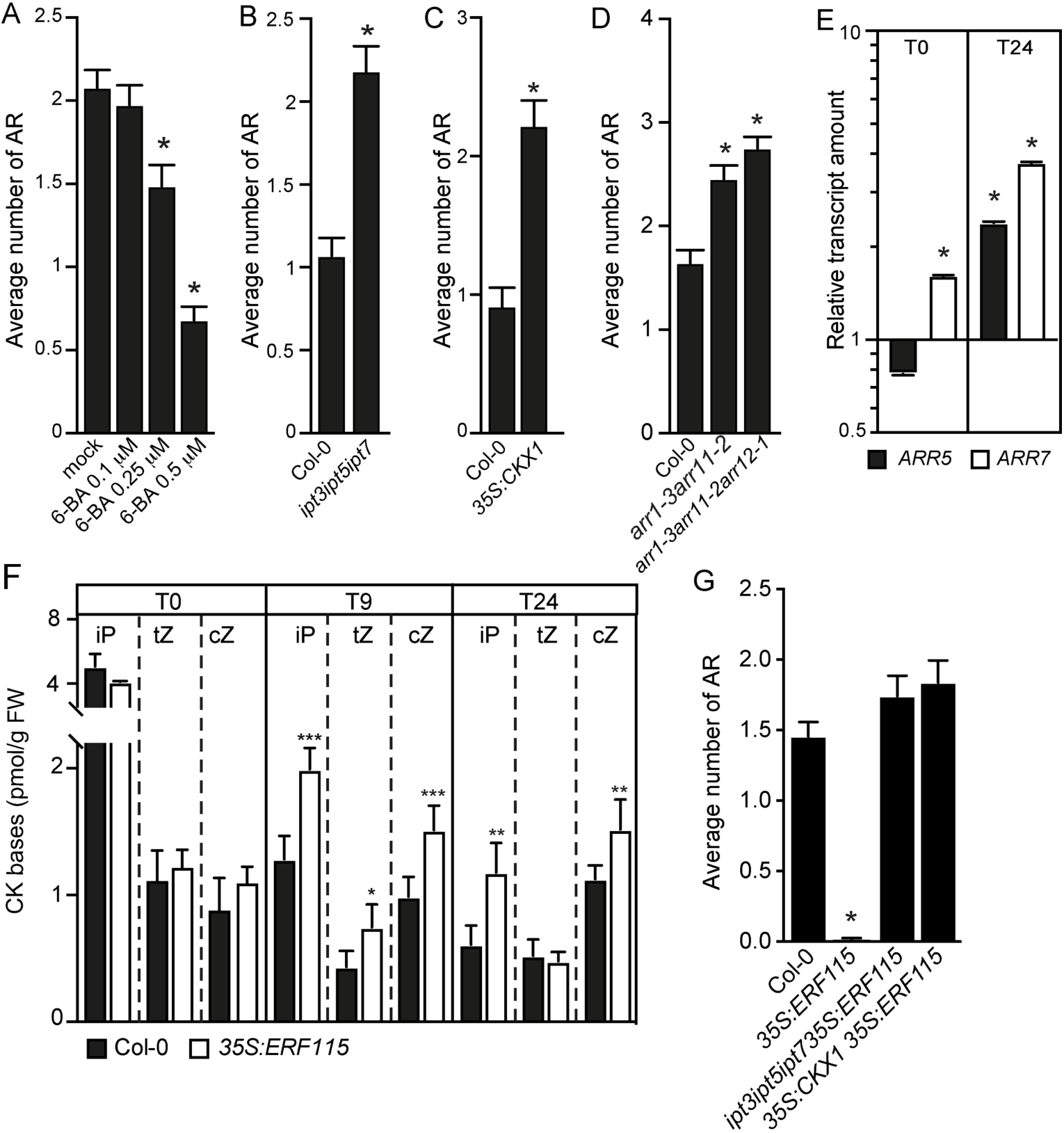
Cytokinins inhibit ARI downstream of *ERF115*. (A) Average numbers of ARs produced by wild-type (Col-0) seedlings, which were grown in the dark until their hypocotyls were 6-7 mm long, then transferred to fresh medium containing either mock solution or solutions with indicated concentrations of 6-Benzylaminopurine (6-BA). The seedlings were kept for seven more days under long-day conditions to induce ARs. Seedlings treated with 0.25 µM or 0.5 µM 6-BA significantly differed from the mock-treated controls, according to a non-parametric Kruskal-Wallis test followed by Dunn’s multiple comparison test. Error bars indicate ± SEM (n ≥ 40, P < 0.004). (B) to (D) Average numbers of ARs produced by wild-type plants and: (B) *ipt3ipt5ipt7* triple mutants defective in CK biosynthesis, (C) *35S:CKX1 CYTOKININ OXIDASE1*-overexpressing plants, which have reduced CK contents due to increased rates of degradation, and (D) CK signaling mutants. (E) Relative amounts of *ARR5* and *ARR7* transcripts quantified by qRT-PCR. Total RNA was extracted from hypocotyls of *35S:ERF115* and the wild-type seedlings grown in AR-inducing conditions, as outlined above, at T0 (at the end of the dark incubation) and T24 (24 hours later). The gene expression values are relative to wild-type values, which were arbitrarily set to 1. The Y axis scale is a log_10_ scale. Error bars indicate ± SEM obtained from three technical replicates. Asterisks indicate values that significantly differ from wild-type values according to a *t*-test (P < 0.001, n = 3). The experiment was repeated once with an independent biological replicate and gave similar results. (F) Endogenous contents of active CK bases. The CK bases were quantified in the hypocotyls of *35S:ERF115* and the wild-type seedlings grown in the dark until they were 6-7 mm long (T0) and after their transfer to the light for 9 h (T9) or 24 h (T24). Asterisks indicate statistically significant differences between mutant and wild-type plants according to ANOVA (*, **, and *** indicate P-values of 0.05 > P > 0.01, 0.01 > P > 0.001, and P < 0.001, respectively). Error bars indicate ± SD of six biological replicates. (G) Average numbers of ARs produced by *35S:ERF115* plants, *35S:ERF115* plants overexpressing *CKX1* from a *35S:CKX1* construct and the *ipt3,5,7* triple mutant overexpressing *ERF115* from a *35S:ERF115* construct. Numbers produced by the multiple mutants significantly differed from numbers produced by *35S:ERF115* plants according to a non-parametric Kruskal-Wallis test followed by Dunn’s multiple comparison test. Error bars indicate ± SEM (n ≥ 40, P < 0.0001).

## DISCUSSION

Plants develop ARs in response to diverse intrinsic and/or extrinsic (stress-induced) cues (Bellini et al., 2014; Steffens and Rasmussen, 2016) that are perceived by competent cells and trigger extensive reprogramming that results in targeted cells acquiring new identities (Bellini et al., 2014; Lakehal and Bellini, 2018). The process has both high fundamental interest and practical importance as adventitious rooting is often a limiting step in clonal propagation. However, very little is known about the mechanism triggering the cell reprogramming that leads to ARI. Fortunately, intact hypocotyl-derived AR provide ideal model systems to unravel the signaling networks involved in this and other *de novo* organogenesis processes. We have previously shown that auxin controls ARI in Arabidopsis hypocotyls by modulating JA homeostasis (Gutierrez et al., 2009; Gutierrez et al., 2012; Lakehal et al., 2019a), but the JA signaling mechanism involved was not clear. Here, we provide detailed genetic and mechanistic insights into the JA signaling involved in ARI. Notably, *ninja-1* and *ninja-2* loss-of-function mutants produce ARs, albeit fewer than wild-type controls, and several lines of evidence indicate that this is possibly due to NINJA-independent repression of MYC-dependent machinery by a subset of JAZ proteins. For example, JAZ5, JAZ6, JAZ7 and JAZ8 can directly recruit TPL through their EAR motifs independently of NINJA (Kagale et al., 2010; Causier et al., 2012; Shyu et al., 2012), while JAZ1, JAZ3 and JAZ9 can directly recruit HISTONE DEACETYLASE6 (HDA6) (Zhu et al., 2011), which participates in repression of various JA-induced genes’ expression (Zhu et al., 2011). In addition, yeast two-hybrid experiments have shown that JAZ7, JAZ8 and JAZ13 do not interact with NINJA (Pauwels et al., 2010; Shyu et al., 2012; Thireault et al., 2015), and the Jasmonate-associated (Jas) domain of JAZ directly binds to the region containing the JAZ-interaction domain (JID) and TAD domains of MYC2, MYC3 or MYC4 (Zhang et al., 2015). Moreover, MED25 (one of 29 subunits of the MEDIATOR complex) interacts with MYC proteins and recruits the RNA polymerase II-dependent transcriptional machinery at MYC-target genes (Chen et al., 2012; An et al., 2017). MED25 directly interacts with the TAD domain of MYCs, raising the possibly that it competes with JAZ proteins for access to the TAD domain(Zhang et al., 2015). All these findings suggest that some JAZ proteins might block transcriptional activities of MYC transcription factors involved in ARI in a NINJA-independent manner. Further research is needed to decipher the JAZ-dependent JA perception machinery involved in ARI. For this, combining mutants with potentially complementary functionalities, and/or potentially informative expression patterns, may be more illuminating than generating higher-order multiple mutants based on phylogenetic relationships.

Our results indicate that MYC-mediated JA signaling inhibits ARI in both NINJA-dependent and -independent manners. Both pathways act synergistically in control of the JA response, as indicated by the much lower numbers of ARs produced by *ninjamyc2-322B* double mutants than the parental lines (*ninja* and *myc2-322B*) and wild-type controls. Therefore, the strong phenotype of *ninjamyc2-322B* double mutants may be due to de-repression of not only MYC2, MYC3 and MYC4, but also other NINJA-bound transcription factors (if any). Interestingly, this de-repression results in constitutively enhanced JA signaling. Accordingly, our transcriptomic analysis revealed that most of the JAZ genes, which are JA response marker genes (Chini et al., 2007), were highly and constitutively upregulated in *ninja-1myc2-322B* plants throughout the covered developmental stages. Our results are consistent with previous report suggesting that MYC2 controls root expansion in NINJA dependent and -independent manners (Gasperini et al., 2015).

For many years JA was regarded as a solely stress-related plant hormone, but more recently JA signaling has been implicated in several organogenesis and regenerative processes (Asahina et al., 2011; Gutierrez et al., 2012; Lakehal et al., 2019a; Zhang et al., 2019; Zhou et al., 2019), and attempts to identify its downstream targets have begun. Although its role in adventitious rooting seems to be species- and context-dependent (Lakehal and Bellini, 2018), our results indicate that the *ERF115* gene is likely one of the targets acting downstream of JA in this process. This conclusion is strongly supported by the recent finding that MYC2 induces expression of *ERF115* by directly binding its promoter (Zhou et al., 2019). The *ERF115* acts redundantly with its closely-related paralogs *ERF113* and *ERF114,* which have also been implicated in several organogenesis and regenerative processes (Heyman et al., 2018). Here we provide evidence that *ERF115*-mediated ARI inhibition involves modulation of the CK machinery. Physiological approaches have shown that CKs inhibit ARI in several plant species and model systems (Lakehal and Bellini, 2018). In this study, we genetically demonstrated that depleting CKs by either blocking their biosynthesis or enhancing their degradation restores the ARI wild-type phenotype in an *ERF115*-overexpressing line, confirming that *ERF115* represses ARI through CK signaling. Interestingly, the *ERF115* promoter contains a cytokinin-responsive motif, and a yeast one-hybrid screen has shown that ARR1 and ARR20 bind to the promoter of *ERF115* (Ikeuchi et al., 2018). Although direct evidence is needed, these data suggest that cytokinin signaling may also control the abundance of *ERF115* transcripts. The role of this feedback loop in adventitious rooting, if any, awaits further investigation.

## MATERIALS AND METHODS

### Plant material

The quadruple mutant *jaz7jaz8jaz10jaz13* (Thireault et al., 2015) and quintuple mutant *jaz1jaz3jaz4jaz9jaz10*(Campos et al., 2016) were provided by G. Howe (Michigan State University, USA). The single mutants *ninja-1*, *ninja-2* (Acosta et al., 2013), and *myc2-322B* as well as the double mutants *ninja-1myc2-322B*, *ninja-2myc2-322B* and *ninja-1atr2D* (Gasperini et al., 2015) were provided by E.E. Farmer (University of Lausanne, Switzerland). The gain of function allele of *MYC3* (*atr2D*) (Smolen et al., 2002) was provided by J. Bender (Brown University, Rhode Island, USA). The single mutant *erf115* (SALK_021981) and transgenic lines *pERF115:ERF115:SRDX*, and *35S:ERF115* (Heyman et al., 2013) were provided by L. De Veylder (VIB, University of Gent, Belgium). The *rap2-6l-1* mutant (SALK_051006) (Che et al., 2006), *arr1-3arr11-2* (N6980) and *arr1-3arr11-2 arr12-1* (N6986) were provided by the Nottingham Arabidopsis Stock Centre. The transgenic line *35S:CKX1* (Werner et al., 2003) and triple mutant *ipt3ipt5ipt7* (Miyawaki et al., 2006) were provided by T. Schmülling (Freie Universität Berlin, Germany). E.E. Farmer and L. De Veylder also respectively provided the reporter lines *pMYC2:GUSplus*, *pNINJA:GUSplus* and *pNINJA:NINJA:mCITRINE/ninja-1* (Gasperini et al., 2015) and *pERF115:GUS* (Heyman et al., 2013).

### CRISPR-Cas9 cloning, transformation and mutant screening

To generate the loss-of-function allele *erf114*C, two guide RNAs (ERF114_F and ERF114_R, see Supplemental Table 2) were designed, as previously described (Lakehal et al., 2019b), to target the *ERF114* gene’s first exon (Supplemental Figure 3). The guide RNAs were then cloned into the binary vector pHEE401E, the resulting construct was transformed into *Escherichia coli* cells, and the positive clones were selected by PCR, then confirmed by sequencing, following previous protocols (Xing et al., 2014; Wang et al., 2015). The *Agrobacterium*-mediated floral dip method (Clough and Bent, 1998) was used to transform the construct into *rap2-6l-1* or *erf115* mutants. T1 seedlings were screened on Arabidopsis growth medium (Lakehal et al., 2019b) containing 50 μg/ml hygromycin and surviving seedlings were genotyped for deletions in *ERF114* using primers listed in Supplemental Table 2. Several independent homozygous and heterozygous T1 lines were identified. Only homozygous *erf114C* and Cas9-free lines, confirmed by examination of T2 individuals and Cas9-construct genotyping (Xing et al., 2014; Wang et al., 2015), were used for further analysis.

### Tissue-specific complementation: cloning, transformation and transgenic line screening

The pEN-L4-pGATA23-R1 and pEN-L4-pXPP-R1 plasmids (De Rybel et al., 2010) (Andersen et al., 2018) were gifts from T. Beeckman (VIB, Gent, Belgium) and J. Vermeer (University of Zurich, Switzerland), respectively. Plasmids carrying coding sequences of the *NINJA* gene, pEN-L1-NINJA(noSTOP)-L2, and reporter protein, pEN-R2-mCITRINE-L3 (Gasperini et al., 2015), were gifts from E.E. Framer (University of Lausanne, Switzerland). To generate promoter:NINJA:CT fusion protein constructs, the pEN-L4-promoter-R1, pEN-L1-NINJA(noSTOP)-L2 and pEN-R2-mCITRINE-L3 were recombined into the pB7m34gw vector using LR clonaseII plus (Invitrogen) according to the manufacturer’s instructions. All the expression vectors were confirmed by colony PCR and sequencing, then transformed into GV3101 *Agrobacterium tumefaciens* cells, which were used to transform *ninja-1myc2-332B* double mutants using the floral dip method (Clough and Bent, 1998). Single-copy, homozygous lines were selected by cultivating representatives of T2 and T3 generations on Arabidopsis medium (Lakehal et al., 2019b) supplemented with 10 µg/ml DL-phosphinothricin (Duchefa biochemie). At least two lines carrying each construct showing the same phenotype were further characterised.

### Growth conditions and root (adventitious and lateral) phenotyping

Previously described adventitious rooting conditions (Sorin et al., 2005; Gutierrez et al., 2009; Gutierrez et al., 2012; Lakehal et al., 2019a) were applied in all the experiments. Seedlings were etiolated in the dark until the hypocotyls were approximatively 6-7 mm long, then were grown in long-day conditions (16 h light 22° C/ 8h dark 17° C cycles, with 130-140 µmol photons/m^2^/sec during light phases and constant 65% relative humidity). After 7 days, numbers of primordia and emerged ARs were counted under a binocular stereomicroscope. Numbers of visible LRs were also counted, and the primary root length was measured using ImageJ software (Schindelin et al., 2012). The LR density was calculated by dividing the number of LR by the primary root length.

### RNA isolation and cDNA synthesis

Total RNA was prepared using a RNAqueous® Total RNA Isolation kit (Ambion™). Portions (4 μg) of the resulting RNA preparations were treated with DNaseI using a DNA*free* Kit (Ambion™) then cDNA was synthesized by reverse transcription using a SuperScript II Reverse transcriptase kit (Invitrogen) with anchored-oligo(dT)_18_ primers, according to the manufacturer’s instructions.

### Quantitative RT-PCR (qRT-PCR) experiments

Transcript levels were assessed by qRT-PCR, in assays with triplicate reaction mixtures (final volume, 20 μL) containing 5 μL of cDNA, 0.5 μM of both forward and reverse primers, and 1× LightCycler 480 SYBR Green I Master (Roche) using a LightCycler 480 instrument (Roche) according to the manufacturer’s instructions. A melting curve analytical step was added to each PCR program. The sequences of primers used for all target genes are presented in Supplemental Table 2. The crossing threshold (CT) values for each sample were acquired with the LightCycler 480 software (Roche) using the second derivative maximum method. All quantifications were repeated with at least two independent biological replicates.

### qRT-PCR data analysis

Reference genes were validated as the most stably expressed genes in our experimental procedures (Gutierrez et al., 2009) using GenNorm software and the most stable two (*TIP41* and *EF1A*) were used to normalize the quantitative qPCR data. The data obtained using both reference genes were similar and only data obtained using *TIP41* are presented here. Relative transcript amounts were calculated as previously described (Gutierrez et al., 2009), and considered significant if fold differences were ≥ 1.5 with *p*-values ≤ 0.05).

### RNA sequencing and transcriptomic analysis

Total RNA was extracted from etiolated hypocotyls grown in darkness at T0, just before exposure of some of the etiolated seedlings to light. Further samples were collected after 9 and 24 h in long-day conditions (T9L and T24L, respectively). In each case three biological replicates were prepared, and the total RNA was treated with DNaseI using a DNA*free* Kit (Ambion™) to remove any contaminating DNA. The RNA’s integrity and quantity were checked using a 2100 Bioanalyzer (Agilent), then it was sequenced by BGI Tech (China) using an Illumina HiSeq 4000 platform. The reads were trimmed with SOAPnuke then clean reads were mapped to the Araport11 reference sequence using Bowtie2 (Langmead and Salzberg, 2012). Gene expression was quantified using RSEM (RNA-Seq by Expectation-Maximization) (Li and Dewey, 2014) and differentially expressed genes (DEGs) between *ninja-1myc2-322B* and wild-type plants at selected time points were detected using NOISeq software (Tarazona et al., 2011) with fold change ≥ 2 and probability 0.8 settings.

FIMO tools were used, via the http://meme-suite.org/tools/fimo web interface, to scan promoters (1 Kb upstream of ATG translation start codons) of the DEGs for G box and G-box-like motifs with a 1E-4 *p*-value setting.

RNA-seq data has been deposited at the European Nucleotide Archive (ENA) - https://www.ebi.ac.uk/ena and will be available using the following accession number PRJEB36195.

### Spatiotemporal gene expression patterns during AR initiation

The spatiotemporal patterns of *NINJA*, *MYC2* and *ERF115* genes’ expression during AR initiation were monitored by GUS-based analysis, as follows. Seedlings expressing *pNINJA:GUSplus*, *pMYC2:GUSplus* or *pERF115:GUS* were grown in AR-inducing conditions as described above, then stained with X-GLCA (Duchefa Biochemie, X1405.1000) as previously described (Sorin et al., 2005). At least 25 seedlings of each genotype sampled at each time point were stained, and one representative seedling of each set was photographed.

### Sample preparation for hormone quantification

Hypocotyls were collected from seedlings grown in AR-inducing conditions as described above. The hypocotyls were quickly dried on tissue paper then frozen in liquid nitrogen. Samples were prepared from six biological replicates.

### Quantification of *cis*-OPDA, JA and JA-Ile

Endogenous levels of jasmonates (*cis*-OPDA, free JA, and JA-Ile) were determined in 20 mg samples, as previously described (Floková et al., 2014).

### Quantification of endogenous cytokinin bases

Cytokinin metabolites were quantified following published methodology (Svačinová et al., 2012; Antoniadi et al., 2015) Briefly, samples (20 mg FW) were homogenized and extracted in 1 ml of modified Bieleski solvent (60% MeOH, 10% HCOOH and 30% H_2_O) together with a cocktail of stable isotope-labelled internal standards (0.25 pmol of CK bases added per sample). The extracts were applied to an Oasis MCX column (30 mg/ml, Waters) conditioned with 1 ml each of 100% MeOH and H_2_O, equilibrated sequentially with 1 ml of 50% (v/v) nitric acid, 1 ml of H_2_O, and 1 ml of 1 M HCOOH, then washed with 1 ml of 1 M HCOOH and 1 ml 100% MeOH. Analytes were then eluted in two steps with 1 ml of 0.35 M aqueous NH_4_OH solution and 2 ml of 0.35 M NH_4_OH in 60% (v/v) MeOH solution, evaporated to dryness *in vacuo* and stored at −20°C. Cytokinin levels were determined by ultra-high performance liquid chromatography-electrospray tandem mass spectrometry (UHPLC-MS/MS) using stable isotope-labelled internal standards as reference compounds (Rittenberg D., 1940). Following separation with an Acquity UPLC^®^ system (Waters, Milford, MA, USA) equipped with an Acquity UPLC BEH Shield RP18 column (150×2.1 mm dimensions, 1.7 μm particles; Waters), the effluent was introduced into the electrospray ion source of a Xevo™ TQ-S MS triple quadrupole mass spectrometer (Waters). Six independent biological replicates of each genotype sampled at each time point were analyzed.

### Confocal Laser Scanning Microscopy (cLSM) analysis

Images of the vasculature in *Arabidopsis* hypocotyls at depths up to 150 µm from the epidermal surface were acquired using a Zeiss LSM880 inverted confocal laser scanning microscope (Carl Zeiss GmbH, Oberkochen, Germany) equipped with a C-Achroplan 32x/0.85 W Corr M27 lens. The seedlings were etiolated in the dark until their hypocotyls were 6-7 mm long then incubated in liquid medium containing 30 µg/ml propidium iodide (PI) as a cell wall counter-stain to identify the cell layers, and observed while still alive, mounted with the same medium. The PI was excited using a 561 nm laser while expressed reporter protein (mCITRINE) was excited with a 488 nm Argon laser, using a MBS 488/561 Main Beam Splitter. PI Fluorescence from PI and the reporter (mCITRINE) were detected to localize expression with a photomultiplier tube (PMT) detector and a GaAsP (gallium arsenide phosphide photomultiplier tube) 32-channels spectral detector (with about two times higher sensitivity than the PMT, enabling detection of even poorly expressed reporters), respectively. 3D projections and orthogonal views were generated using FIJI/Image J (Schindelin et al., 2012), including image-wide adjustments of brightness and contrast for each channel before merging to ensure that both signals from PI and the fluorescent protein reporter could be easily seen in all displayed images.

## FUNDING

This work was supported by grants from the Swedish Research Council (VR), the Swedish Research Council for Research and Innovation for Sustainable Growth (VINNOVA), the K & A Wallenberg Foundation, the Carl Trygger Foundation, and the Carl Kempe Foundation awarded to C.B, together with grants from the Ministry of Education, Youth and Sports of the Czech Republic (European Regional Development Fund-Project “Plants as a tool for sustainable global development” No. CZ.02.1.01/0.0/0.0/16_019/0000827), and the Czech Science Foundation (Project No. 19-00973S) awarded to O.N.

## ACKNOWLEDGMENTS

The authors sincerely thank Alok Ranjan for his technical support, Hana Martínková and Petra Amakorová for their help with phytohormone analyses, Nicolas Delhomme and Iryna Shutava from the UPSC bioinformatic platform for their advice and help with the RNA-Seq data analysis and *cis-*regulatory motif search.

## AUTHORS’ CONTRIBUTIONS

Methodology, A.L., C.B., A.D. Investigation, A.L., A.D., Z.R., S.A., S.E., O.N. Writing – original draft, A.L., A.D. Writing –Review & Editing, A.L., C.B.; Conceptualization, A.L. C.B.; Supervision, C.B.; Funding Acquisition, C.B., H.T., O.N. and M.S.

## Notes

#### Summary of Updates

Added URL and accession number for RNAseq data Modified Acknowledgements

https://www.ebi.ac.uk/ena

