## Supplemental information for "*ETHYLENE RESPONSE FACTOR 115* integrates jasmonate and cytokinin signaling machineries to repress adventitious rooting in *Arabidopsis*"

**This PDF includes:**

Supplemental Figures 1 to 5

Supplemental Tables 2

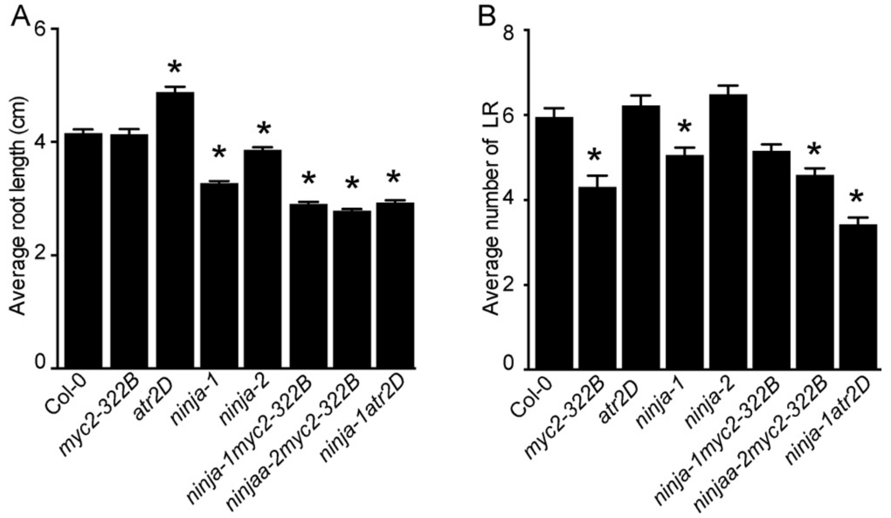


**Supplemental Figure 1 Jasmonate signaling affects primary root (PR) length and lateral root (LR) number.**

(A) PR lengths of wild-type (Col-0) plants and JA signaling mutants grown in AR phenotyping conditions. Asterisks indicate significant differences between the mutants and wild-type plants according to one-way ANOVA followed by Tukey's multiple comparison test. Error bars indicate ± SEM ($n \geq 25$, $P < 0.05$).

(B) Numbers of LRs produced by wild-type plants and JA signaling mutants grown in AR phenotyping conditions. Asterisks indicate significant differences between the mutants and wild-type plants according to one-way ANOVA followed by Tukey's multiple comparison test. Error bars indicate ± SEM ($n \geq 25$, $P <0.05$).

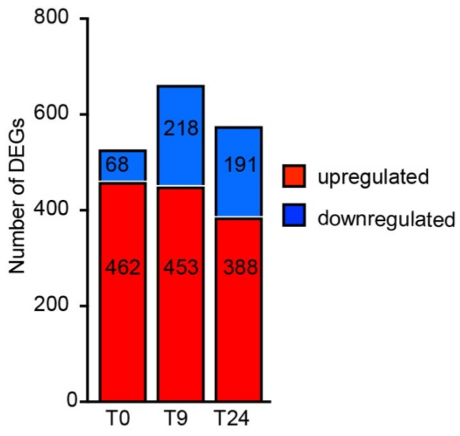


**Supplemental Figure 2 Numbers of DEGs detected in RNA-Seq experiments.**

Red and blue colors respectively indicate up- and down-regulated genes in the *ninja-1myc2-322B* double mutant relative to wild-type expression levels.

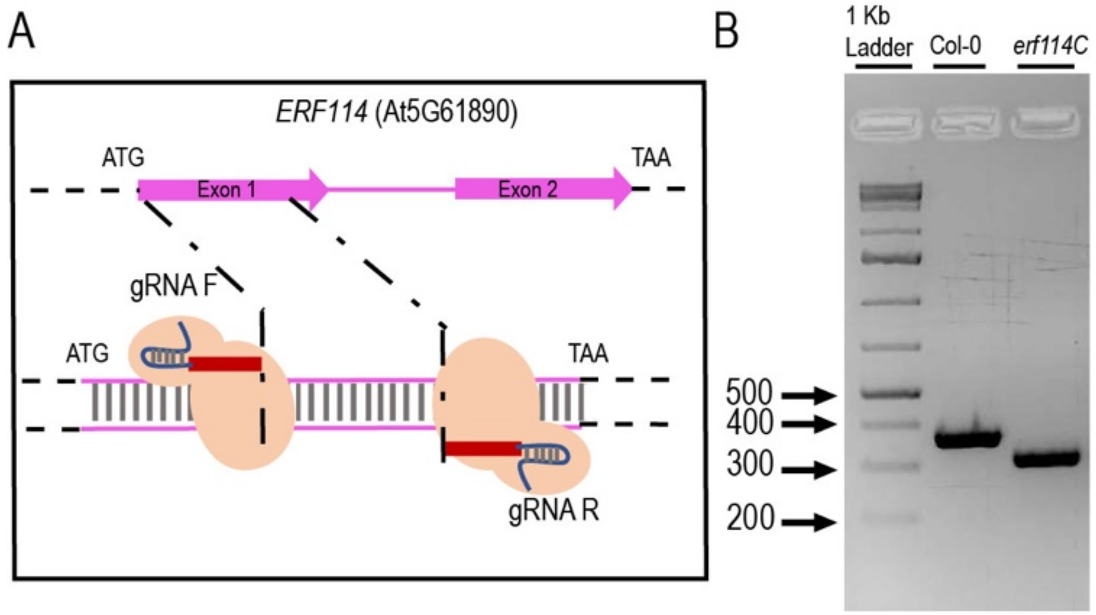


**Supplemental Figure 3 Illustration of the CRISPR-Cas9 strategy**.

(A) Two guide RNAs were designed to target a relatively large DNA fragment of the *ERF114* gene in a *rap2-6l-1* or *erf115* loss-of-function mutant background.

(B) Photo of a representative agarose gel indicating sizes of the wild-type allele *ERF114* and variant *erf114C* with a deletion.

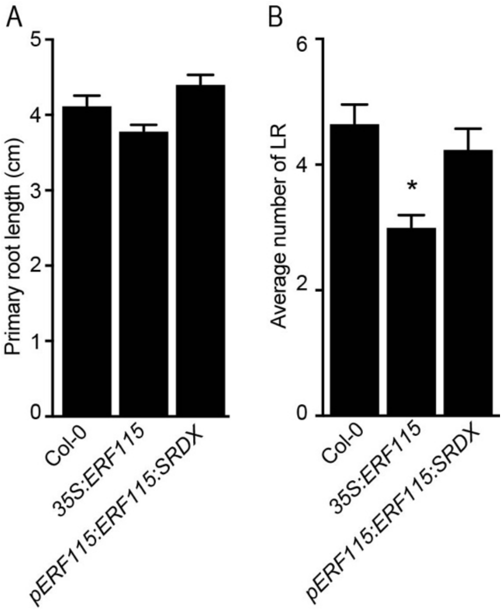


**Supplemental Figure 4 Overexpressing *ERF115* affects the lateral root number.**

(A) Primary root lengths of wild-type, *35S:ERF115* and *pERF115:ERF115:SRDX* plants grown in AR phenotyping conditions. A one-way ANOVA followed by Tukey's multiple comparison post-test. Error bars indicate ± SEM ($n \geq 25$, $P < 0.06$).

(B) Numbers of lateral roots produced by wild-type, *35S:ERF115* and *pERF115:ERF115:SRDX* plants grown in AR phenotyping conditions. The asterisk indicates a significant difference between *35S:ERF115* and wild-type plants according to one-way ANOVA followed by Tukey's multiple comparison post-test. Error bars indicate ± SEM ($n \geq 25$, $P < 0.0001$).

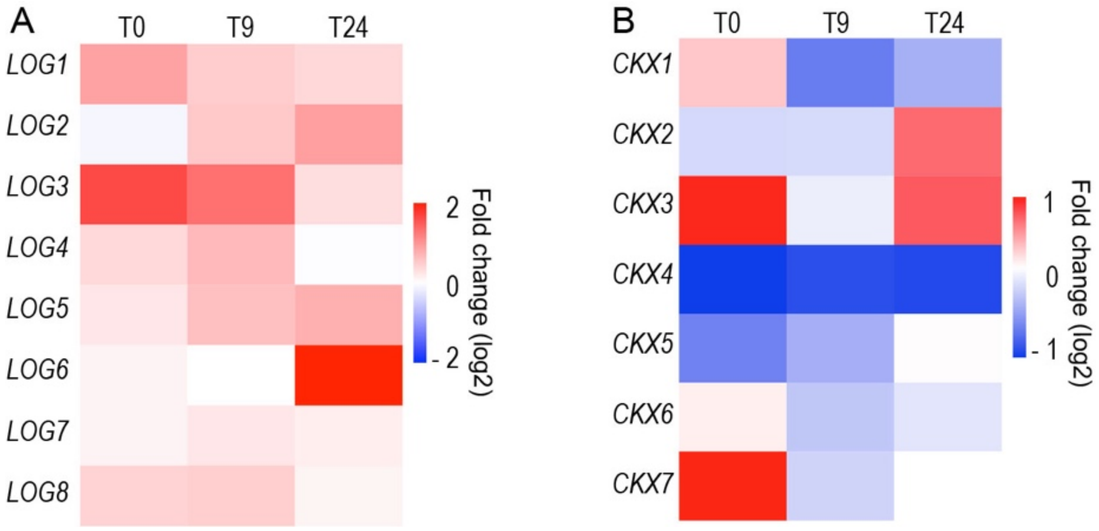


**Supplemental Figure 5 Heatmaps of expression of (A) LOG genes and (B) CKX genes.** The map is based on fold-differences ($\log_2$) in transcript abundance (based on RNA-seq data) in *ninja-1myc2-322B* double mutant samples relative to wild-type samples. Red and blue colors respectively indicate up- and down-regulated genes in *ninja-1myc2-322B* double mutant relative to wild type expression levels.

### Supplemental Table 2: Primers used for qRT-PCR, cloning and genotyping

| Primer name | Gene number | Forward primer | Reverse primer |
|---|---|---|---|
| *qRT-ERF113* | AT5G13330 | CAAGGCCCTACTACCAC CACAA | GGTCGAGGAGGAGGTGAGTTC |
| *qRT-ERF114* | AT5G61890 | AGAACTTGTTCCCGGTC TTCTCG | AGTCAAGGCCGAGACCATAACAC |
| *qRT-ERF115* | AT5G07310 | GGAAACCAAAGCAGCTCTCA | GCAGCTTCAGCAGTCTCAAA |
| *qRT-ARR5* | AT3G48100 | TGTCGATAGTGCGACAAGAGC | CTTCAGATCCTCAAATCCAACC |
| *qRT-ARR7* | AT1G19050 | TCAATGCCAGGACTTTCAGGA | TGCTCCTTCTTTGAGACATTCTTG |
| *qRT-TIP41* | At4g34270 | GCTCATCGGTACGCTCTTTT | TCCATCAGTCAGAGGCTTCC |
| *qRT-EF1*A | At5g60390 | TGGTGACGCTGGTATGGTTA | TCCTTCTTGTCCACGCTCTT |
| *Geno-ERF114* | AT5G61890 | GATGTTCAACGATGCCATAAAGA | GATGAGTAGGCGCAACTTGTT |
| *gRNA_ERF114* | AT5G61890 | GAGATCGGGCCGAGAAGAC | ATTACTTGAGTCAAGGCCG |
